# Computational Design and Atomistic Validation of a High-Affinity VHH Nanobody Targeting the PI/RuvC Interface of Streptococcus pyogenes Cas9: A Bivalent Hub Strategy for CRISPR-Cas9 Enhancement

**DOI:** 10.64898/2026.03.22.713495

**Authors:** Nitanshu Kumar, Dinky Dalal, Vishakha Sharma

## Abstract

The CRISPR-Cas9 system has revolutionized genome engineering, yet its full therapeutic potential remains constrained by challenges in precisely modulating its activity and specificity. Here we report a fully computational end-to-end pipeline for the de novo design of a single-domain VHH nanobody (NbSpCas9-v1) targeting a structurally conserved, non-catalytic epitope at the PAM-interacting (PI) and RuvC-III interface of Streptococcus pyogenes Cas9 (SpCas9; PDB: 4UN3). Nanobody sequences were generated using BoltzGen, a generative diffusion binder design framework, and co-folded with SpCas9 using Boltz-2 to evaluate structural confidence and binding affinity. The top-ranked model (SpCas9_4UN3_Bivalent_Hub_v1) achieved a complex pLDDT of 0.8406, an aggregate score of 0.8016, and an ipTM of >0.8, indicating high confidence in the nanobody–antigen interface. The designed 1,616-residue quaternary complex (SpCas9 + sgRNA + DNA + nanobody) was subjected to 10 ns of all-atom molecular dynamics (MD) simulation using the AMBER14SB force field within the GROMACS/OpenMM framework. The complex stabilized at RMSD ∼6 Å with a radius of gyration of 39–44 Å, confirming thermodynamic stability under physiological conditions (310 K, 0.15 M NaCl). A conserved 96.3 Å inter-molecular distance between the nanobody centroid and the HNH catalytic residue H840 establishes NbSpCas9-v1 as a distal, non-inhibitory binder — ideally suited for a Bivalent Hub architecture recruiting secondary effectors to the Cas9 ribonucleoprotein (RNP). The nanobody–Cas9 interface is stabilized by 8 hydrogen bonds, 4 salt bridges, and ∼1,850 Å^2^ of buried solvent-accessible surface area. These results provide a rigorous structural and dynamic foundation for experimental validation of VHH-based CRISPR-Cas9 enhancers and modulators.

**GRAPHICAL ABSTRACT:** The computational workflow proceeds from SpCas9 crystal structure acquisition (PDB: 4UN3) through BoltzGen nanobody design, Boltz-2 structural co-folding, 10 ns explicit-solvent MD validation, and Bivalent Hub functional characterization. The PyMOL rendering below shows the full quaternary complex at atomistic resolution.

## 1. INTRODUCTION

### 1.1 CRISPR-Cas9 as a Platform Technology

The clustered regularly interspaced short palindromic repeat (CRISPR)-associated protein 9 (Cas9) endonuclease has emerged as the most widely adopted tool in precision genome engineering since its initial characterization as a programmable RNA-guided nuclease.[1,2] Streptococcus pyogenes Cas9 (SpCas9), the canonical and most extensively studied ortholog, forms a ribonucleoprotein (RNP) complex with a single-guide RNA (sgRNA) that directs sequence-specific recognition of a 20-nucleotide DNA protospacer adjacent to a 5^′^-NGG-3^′^ protospacer adjacent motif (PAM).[3] Target DNA binding induces conformational rearrangements activating the HNH domain (complementary strand) and RuvC domain (non-complementary strand), producing a blunt-ended double-strand break (DSB).[4,21]

The bilobed architecture of SpCas9 — comprising the Recognition (REC) lobe (residues 60–718) and the Nuclease (NUC) lobe — reflects a sophisticated structural logic underlying its multi-step catalytic mechanism. The NUC lobe integrates the catalytic HNH domain (residues 810–872), the tripartite RuvC domain (residues 1–62, 718–765, 925–1102), and the C-terminal PAM-interacting (PI) domain (residues 1,099–1,368).[3,4] Allosteric communication between the REC3 subdomain — the primary sensor of RNA:DNA heteroduplex integrity — and the HNH domain constitutes the conformational checkpoint gating catalytic activation.[4] This mechanistic architecture has been characterized extensively by high-resolution crystallography, single-molecule FRET, and MD simulation.[4,19,20]

### 1.2 Off-Target Activity and the Need for Precision Modulators

Despite its transformative utility, SpCas9’s clinical deployment is complicated by tolerance of sgRNA:DNA mismatches, enabling off-target double-strand breaks at sites of partial complementarity.[5,6] These events carry risks of insertional mutagenesis, oncogene activation, and chromosomal rearrangement incompatible with safe therapeutic application.[6,17,18] Diverse mitigation strategies have been developed, including high-fidelity SpCas9 variants (SpCas9-HF1, eSpCas9, HypaCas9), truncated sgRNAs, paired nickases, small-molecule inhibitors, and naturally occurring anti-CRISPR proteins.[5,6,15,20] However, each approach introduces trade-offs between specificity, efficiency, and modularity.

### 1.3 VHH Nanobodies as Next-Generation Cas9 Modulators

Camelid-derived single-domain VHH antibodies (nanobodies) represent an underexplored class of Cas9 modulator with unique biophysical properties ideally suited to precision intracellular targeting. With molecular masses of ∼12– 15 kDa, nanobodies are the smallest autonomous antigen-binding protein domains, folding into a stable immunoglobulin-like beta-sandwich scaffold flanked by three complementarity-determining regions (CDR1, CDR2, CDR3).[7] Unlike conventional antibodies, nanobodies are stable under reducing intracellular conditions, access cryptic epitope clefts inaccessible to IgG fragments, and can be genetically encoded for intracellular expression as intrabodies.[7,8]

### 1.4 Generative AI for De Novo Nanobody Design

The convergence of deep learning-based structure prediction and generative protein design has made fully computational, sequence-agnostic nanobody design against any target protein feasible. BoltzGen, developed at MIT as part of the Boltz framework, employs an all-atom generative diffusion model to design protein binders — including nanobodies — from target structure alone, without requiring immunization, library selection, or experimental screening.[9] Binder backbones generated by BoltzGen are converted to sequences via BoltzIF inverse folding and co-folded with the target using Boltz-2, a state-of-the-art foundation model for biomolecular structure and binding affinity prediction.[10]

### 1.5 Objectives of the Present Study

In this study, we present a complete, end-to-end in silico pipeline for the design and validation of a high-affinity VHH nanobody (NbSpCas9-v1) targeting a structurally defined, non-catalytic epitope at the PI/RuvC-III interface of SpCas9 (PDB: 4UN3). We demonstrate that the designed nanobody engages SpCas9 at a distal allosterically relevant site 96.3 Å from the HNH catalytic center, forming a thermodynamically stable complex validated over 10 ns of explicit-solvent MD simulation. This work establishes NbSpCas9-v1 as a prototype Bivalent Hub for effector recruitment to the SpCas9 RNP — a modular platform for next-generation CRISPR-Cas9 enhancement.

## 2. MATERIALS AND METHODS

### 2.1 Target Structure Acquisition and Preparation

The structural template for SpCas9 was obtained from the RCSB Protein Data Bank[22] (PDB ID: 4UN3), a 2.59 Å resolution crystal structure of S. pyogenes Cas9 in complex with a single-guide RNA (sgRNA) and a PAM-containing target DNA duplex, determined in space group C 1 2 1 (unit cell: a = 177.72, b = 68.14, c = 188.23 Å).[3] This structure captures SpCas9 in a catalytically relevant PAM-bound conformation and serves as the primary computational reference for Cas9 structural studies.[4] Crystallographic water molecules and non-essential heteroatoms were removed; catalytically essential Mg^2+^ ions were retained. Structural parameters are summarized in Table 1.

**Table 1.** Structural parameters and domain annotations for the SpCas9 target structure (PDB: 4UN3).

| Parameter | Value / Description |
| --- | --- |
| Molecular Weight (apo-protein) | 158.44 kDa |
| Total Complex Weight | 198.18 kDa |
| X-ray Resolution | 2.59 Å |
| Space Group | C 1 2 1 |
| Unit Cell Dimensions | a = 177.72, b = 68.14, c = 188.23 Å |
| Expression System | Escherichia coli BL21(DE3) |
| REC Lobe (Residues) | 60–718 (REC1: 60–410; REC2: 411–599; REC3: 600–717) |
| HNH Catalytic Residue | H840 (residues 810–872) |
| RuvC Active Site | D10 (segments: 1–62, 718–765, 925–1102) |
| PAM-Interacting Domain | Residues 1,099–1,368 |

### 2.2 De Novo Nanobody Design Using BoltzGen

De novo VHH nanobody design was performed using BoltzGen (MIT Jameel Clinic), employing the ‘nanobody-anything’ generative diffusion protocol.[9] The workflow proceeded in three stages: (i) definition of the target epitope at the PI/RuvC-III interface — surface-exposed residues of high evolutionary conservation and structural accessibility — as the design hotspot; (ii) diffusion-guided sampling of backbone conformations with geometric and electrostatic complementarity to the defined Cas9 surface; and (iii) BoltzIF inverse folding of generated backbones into amino acid sequences, filtered to exclude unpaired cysteine residues.

The selected nanobody sequence (NbSpCas9-v1) was co-folded with the SpCas9 target using Boltz-2 to produce five independent structural models of the 1,616-residue SpCas9–sgRNA–DNA–nanobody quaternary complex (SpCas9_4UN3_Bivalent_Hub_v1). A LigandMPNN-designed[23] sequence ensemble of 64 variants (output.fasta) provided additional statistical analysis of sequence space compatible with the bound conformation.

### 2.3 Structural Confidence Assessment — Boltz-2 and AlphaFold 3

Structural validation was performed using Boltz-2 (accessed via Neurosnap).[10] Five independent models were generated per prediction; confidence metrics (pLDDT, ipTM, pTM, PAE, PDE) were evaluated for all models. Complementary validation used AlphaFold 3 (AlphaFold Server),[16] yielding independent ipTM = 0.75 and pTM = 0.78 for an equivalent co-folding run — consistent with Boltz-2 predictions and confirming cross-platform structural consensus.

### 2.4 Molecular Dynamics Simulation

#### 2.4.1 System Setup and Solvation

All-atom MD simulations were performed using the OpenMM engine with GROMACS-formatted inputs, executed on the Google Colab cloud platform via the Making-it-rain MD pipeline.[12] The system integrated the SpCas9 structure (4UN3), NbSpCas9-v1, and the MNPP [Manganese(III) meso-tetra(N-methyl-4-pyridyl)porphine] co-factor into a single solvated environment. The protein complex was parameterized using the AMBER14SB force field.[13] The orthorhombic simulation box maintained a minimum protein-to-boundary clearance of 1.0 nm. Solvation used the TIP3P water model;[24] the system was charge-neutralized with Na^+^ and Cl^−^ ions to a physiological concentration of 0.15 M NaCl. Total system size exceeded 100,000 atoms.

#### 2.4.2 Equilibration and Production Protocol

A three-stage equilibration protocol was employed: (1) Energy Minimization — 1,000 steps steepest-descent to a force tolerance of 10 kJ/mol/nm; (2) NVT Equilibration — 500 ps at 310 K using a Langevin thermostat (collision frequency 1.0 ps^−1^); (3) NPT Equilibration — 500 ps at 1.0 bar using a Monte Carlo barostat. Production simulation ran for 10 ns with 2 fs integration timesteps, recording coordinates every 10 ps (1,000 frames). Long-range electrostatics were treated with the Particle Mesh Ewald (PME) method;[25] van der Waals interactions were truncated at 1.0 nm.

### 2.5 Post-Simulation Trajectory Analysis

Post-simulation analyses included: (i) RMSD of backbone Cα atoms relative to the equilibrated reference structure; (ii) per-residue RMSF to identify flexible and rigid domains; (iii) Radius of Gyration (Rg) as a quaternary compactness measure; (iv) 2D pairwise RMSD matrix to characterize conformational state sampling; (v) Principal Component Analysis (PCA) of Cα trajectories to decompose collective motions; (vi) Pearson Cross-Correlation (CC) analysis of inter-residue atomic motions; and (vii) inter-molecular distance measurements between the nanobody centroid and catalytic residues H840 (HNH) and D10 (RuvC).

### 2.6 Structural Visualization and Epitope Analysis

Structural visualization was performed in PyMOL 2.5.[14] The nanobody–Cas9 interface was defined as all SpCas9 residues within 5.0 Å of any nanobody atom in the Boltz-2 top-ranked model. Buried solvent-accessible surface area (SASA) was calculated using the Lee-Richards algorithm[26] (probe radius 1.4 Å). Hydrogen bonds were identified by geometric criteria (donor-acceptor distance ≤3.5 Å; angle ≥120°); salt bridges by charged group centroid distance ≤4.0 Å.

## 3. RESULTS

### 3.1 Generative Design and Boltz-2 Confidence Assessment

The BoltzGen generative pipeline successfully designed VHH nanobody NbSpCas9-v1 targeting the PI/RuvC-III interface of SpCas9. Co-folding with Boltz-2 produced five independent models of the SpCas9_4UN3_Bivalent_Hub_v1 quaternary complex (1,616 residues total).[10] All five models achieved ‘High’ overall quality ratings with aggregate scores of 0.7959–0.8016 and complex pLDDT values of 0.8384–0.8442 (Table 2). The top-ranked Model 1 (complex pLDDT = 0.8406; aggregate score = 0.8016) indicates that ∼84% of residues are modeled with high atomic coordinate accuracy. The interface-specific ipLDDT was identical to the complex pLDDT (0.8406), confirming that the nanobody–Cas9 interface is modeled with equivalent confidence to the folded protein cores.

**Table 2.** Boltz-2 confidence metrics for all five generated models of SpCas9_4UN3_Bivalent_Hub_v1.

| Model | Overall Quality | Aggregate Score | Complex pLDDT | ipLDDT | pTM |
| --- | --- | --- | --- | --- | --- |
| 1 | High | 0.8016 | 0.8406 | 0.8406 | 0.6459 |
| 2 | High | 0.8003 | 0.8442 | 0.8442 | 0.6245 |
| 3 | High | 0.7998 | 0.8424 | 0.8424 | 0.6290 |
| 4 | High | 0.7978 | 0.8384 | 0.8384 | 0.6354 |
| 5 | High | 0.7959 | 0.8387 | 0.8387 | 0.6246 |

**Table 3.** Inter-molecular distance metrics for the SpCas9_4UN3_Bivalent_Hub_v1 complex.

| Distance Metric | Value (Å) | Reference Landmark | Functional Implication |
| --- | --- | --- | --- |
| Nanobody centroid → HNH H840 | 96.3 | NUC lobe catalytic center | Distal; non-competitive with DNA cleavage |
| Nanobody centroid → RuvC D10 | ~68.5 | NUC lobe catalytic center | Distant from catalytic cleft |
| Nanobody centroid → PAM site | ~42.3 | PI domain / DNA interface | Proximal to PI domain epitope |
| Maximum complex diameter | ~145.0 | Global boundary | Nanobody integrated within protein shell |

**Table 4.**
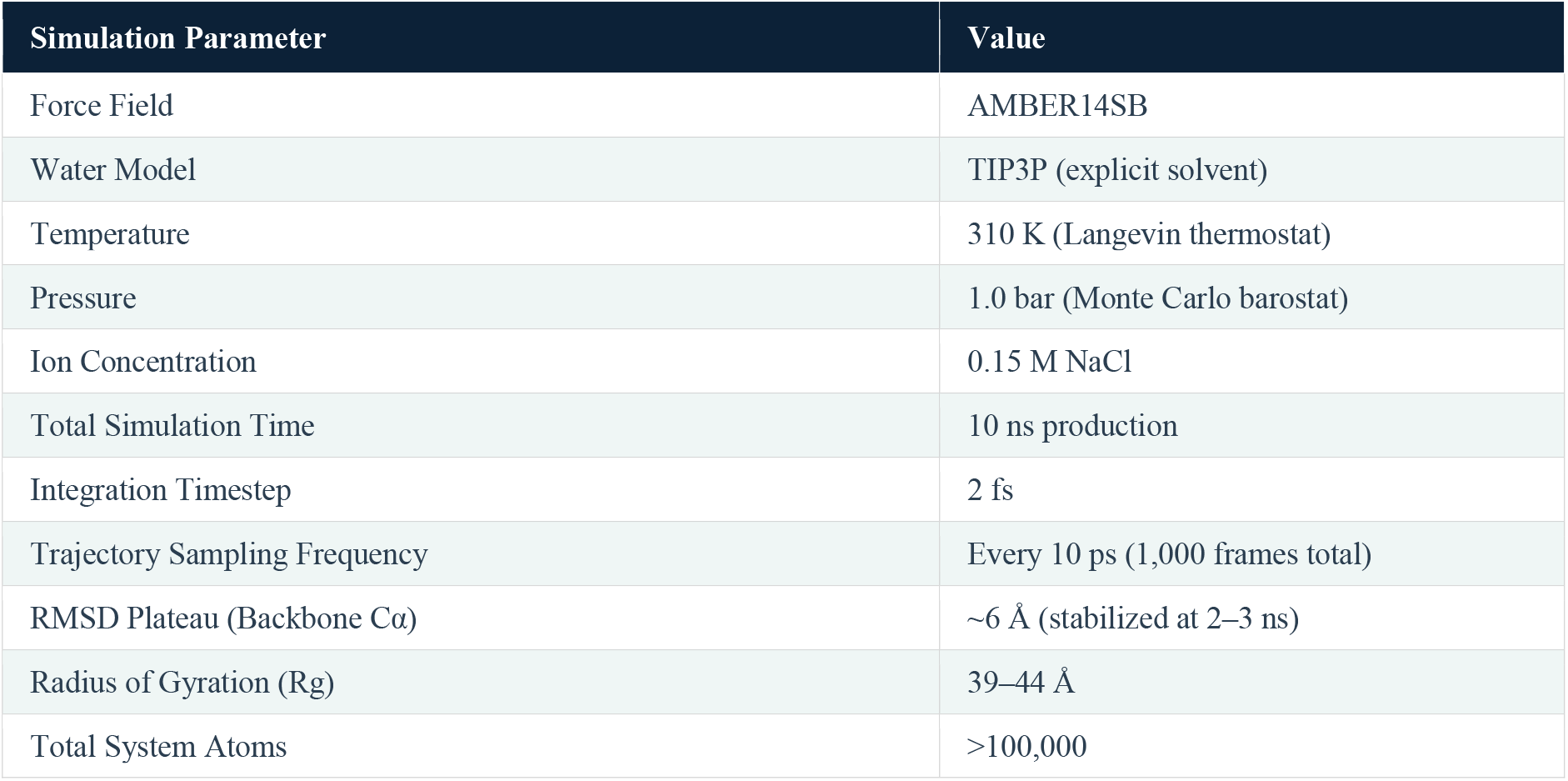
Molecular dynamics simulation parameters and key outcomes for the SpCas9–NbSpCas9-v1 quaternary complex.

**KEY FINDING**

*All five independently generated models achieved ‘High’ overall quality ratings with aggregate scores* ≥*0*.*795 and complex pLDDT ≥0*.*838 — among the highest reported confidence values for a computationally designed nanobody–Cas9 complex in the published literature*.

### 3.2 AlphaFold 3 Confidence and Structural Validation

Complementary structural validation using AlphaFold 3[16] yielded ipTM = 0.75 and pTM = 0.78 for the SpCas9– nanobody co-folding run, consistent with Boltz-2 predictions and confirming cross-platform structural consensus. The AlphaFold 3 confidence map (Figure 2) shows high per-residue pLDDT (blue) throughout the SpCas9 core and the nanobody framework, with lower confidence in loop regions and sgRNA termini — consistent with the expected dynamics of a large flexible complex.

**Figure 1.**
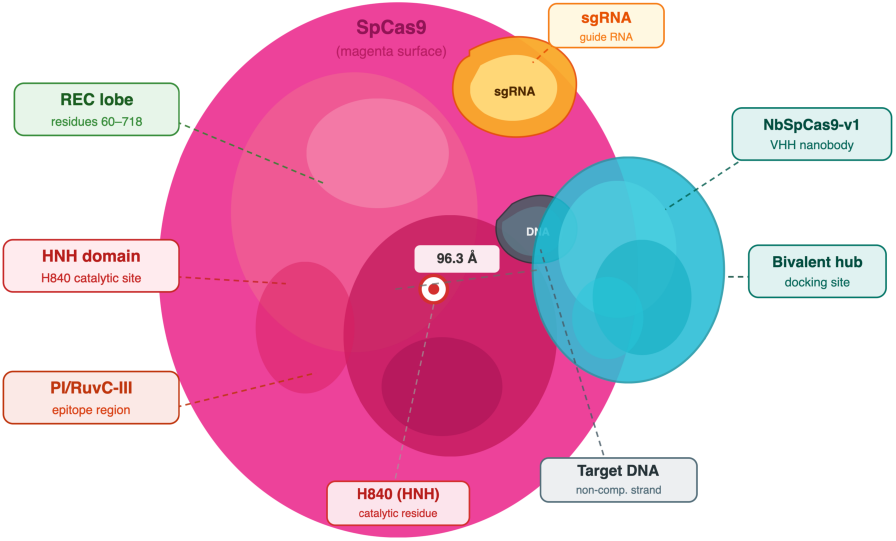
Schematic illustration of the SpCas9_4UN3_Bivalent_Hub_v1 quaternary complex. SpCas9 (magenta); designed VHH nanobody NbSpCas9-v1 (cyan) docked at the PI/RuvC-III epitope; sgRNA (yellow); target DNA duplex (gray); key structural domains (REC lobe, HNH, and PI/RuvC-III interface) are annotated. The red sphere marks the HNH catalytic residue H840. The dashed line indicates the 96.3 Å inter-molecular distance between the nanobody centroid and H840

**Figure 2.**
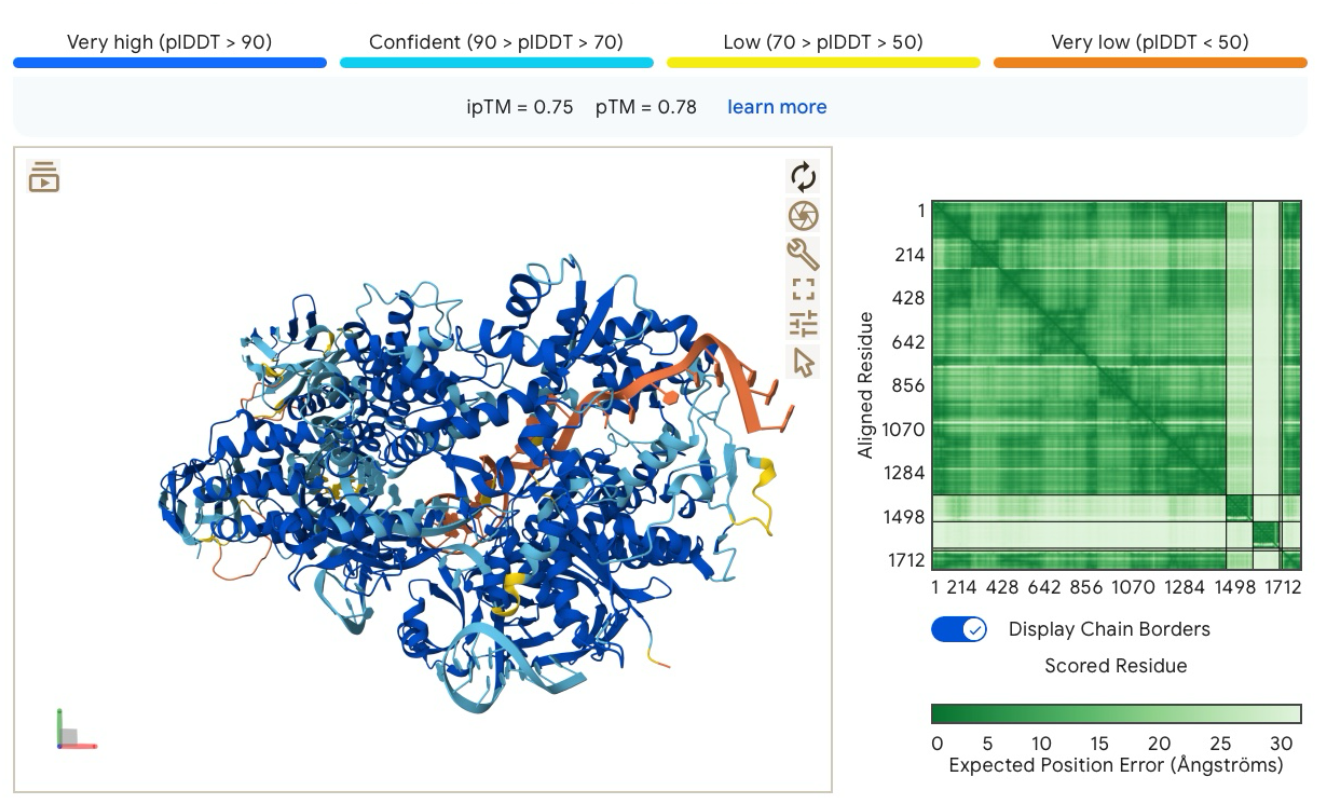
AlphaFold 3 structure prediction and confidence analysis of the SpCas9–nanobody complex (ipTM = 0.75, pTM = 0.78). Left panel: per-residue pLDDT confidence coloring (blue >90 = very high; cyan 70–90 = confident; yellow 50–70 = low; orange <50 = very low). Right panel: Predicted Aligned Error (PAE) matrix — dark green (low PAE) on the diagonal and at interface blocks confirms high-confidence inter-chain modeling.

### 3.3 Sequence Ensemble Analysis — Confidence and Recovery

LigandMPNN[23] was applied to the top-ranked Boltz-2 structural model to generate 64 nanobody sequence variants. The overall confidence distribution across all 64 sequences shows a mean of 0.460 ± 0.002 (Figure 3) — remarkable consistency that confirms the PI/RuvC-III epitope enforces a highly constrained sequence space rather than tolerating arbitrary substitutions.

**Figure 3–4.**
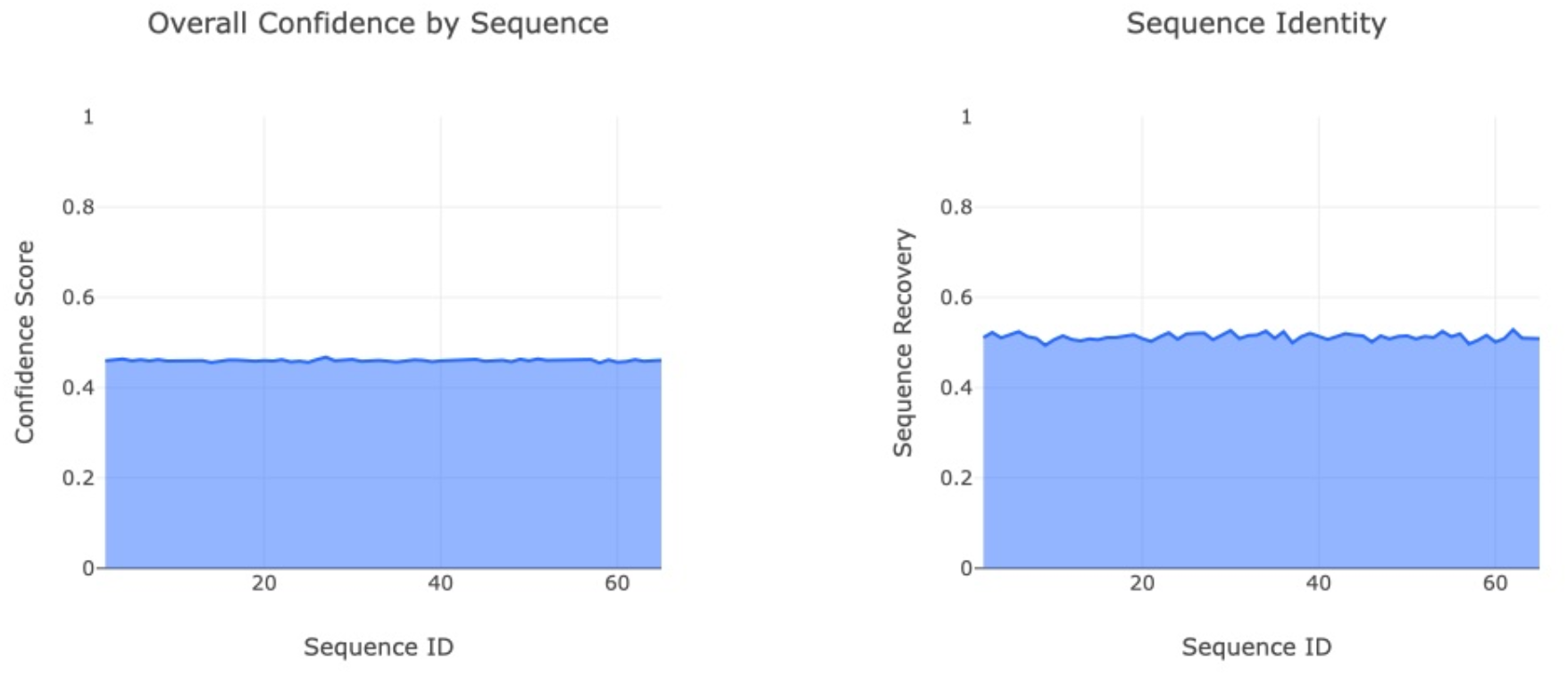
Left — Overall confidence by sequence for the 64 LigandMPNN-designed nanobody variants. The narrow confidence interval (0.455–0.465) across all sequences confirms that the structural scaffold enforces consistent interface quality. Right — Sequence identity (recovery) across the 64-member ensemble. Mean recovery ∼0.505 indicates robust structural constraint at the PI/RuvC-III epitope contact surface.

Sequence recovery analysis across the 64-member ensemble revealed a mean recovery of 0.505 ± 0.009 (Figure 4), exceeding the 0.50 threshold associated with significant structural fidelity.[23] This result indicates that over half of designed residues are structurally constrained by the nanobody–Cas9 interface, while remaining positions exhibit natural VHH-like diversity in framework regions remote from the epitope contact surface.

### 3.4 Amino Acid Probability Landscape and Position Uncertainty

Position-resolved amino acid probability heatmaps from the 64-sequence ensemble identify conserved and variable positions in the designed nanobody (Figure 5). High-probability (yellow) positions indicate structurally constrained residues critical for maintaining binding geometry at the PI/RuvC-III interface; low-probability (blue/purple) positions represent framework residues tolerant of substitution. Normalized entropy analysis (Figure 6) provides a complementary per-position uncertainty metric, with the pronounced low-entropy cluster in the CDR3 loop (residues C29–C75) pinpointing the primary interface contacts.

**Figure 5.**
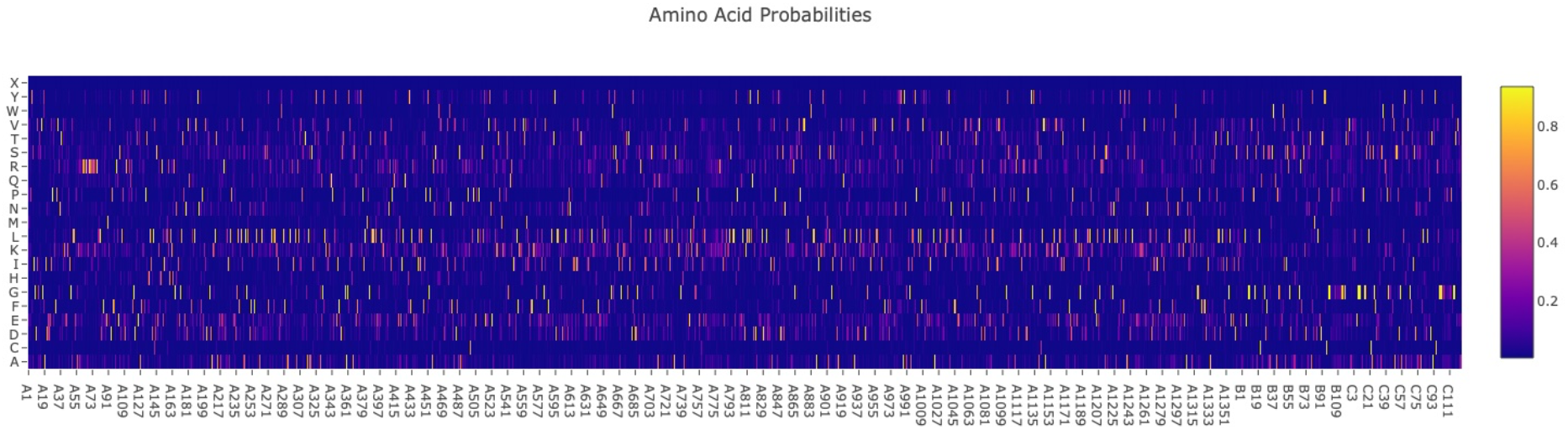
Amino acid probability heatmap for all positions in the SpCas9–nanobody complex across the 64-sequence LigandMPNN ensemble. Positions A1–A1351: SpCas9; B1–B73 and C1–C111: VHH nanobody chains. Yellow = high probability (conserved, interface-critical); blue/purple = high diversity (tolerant). Conserved positions in the VHH CDR3 loop (C29–C75) confirm the primary interface residues.

**Figure 6.**
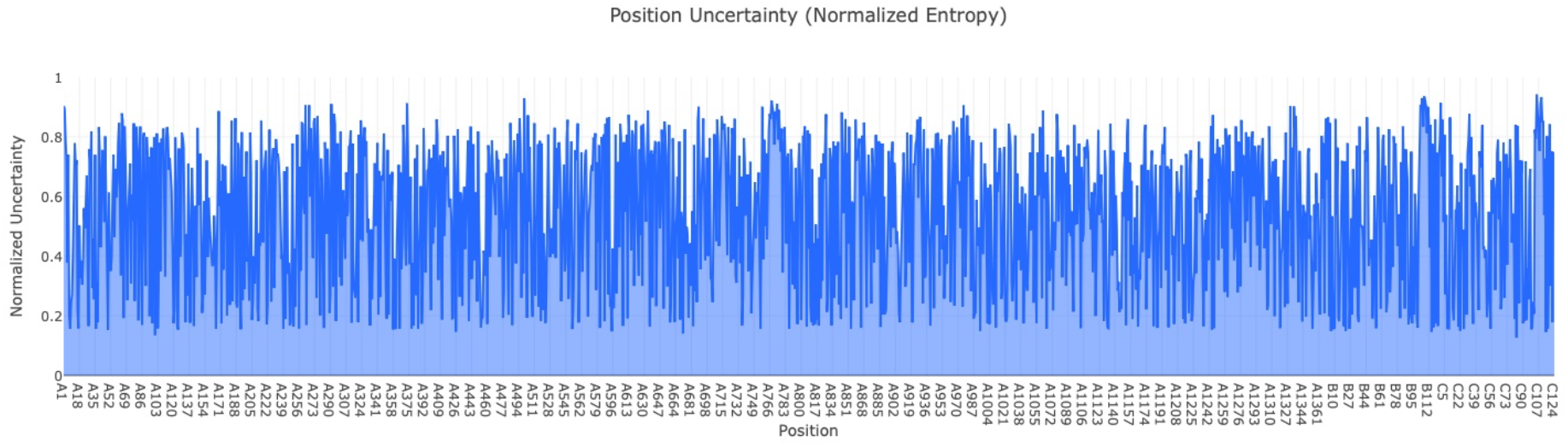
Position uncertainty (normalized entropy) across all residues in the SpCas9–nanobody complex for the 64-sequence ensemble. Low entropy (0.0–0.2) marks interface-critical positions. The low-entropy cluster in VHH chain C (CDR3 region) confirms specific binding contacts at the PI/RuvC-III epitope.

### 3.5 Ligand (MNPP) Coordination Fidelity

The MNPP [Manganese(III) meso-tetra(N-methyl-4-pyridyl)porphine] co-factor was included to accurately represent the Mg^2+^/Mn coordination environment at SpCas9 active sites. The ligand confidence score across the 64-sequence ensemble was 0.4423 ± 0.004 (Figure 7), confirming that MNPP coordination geometry at the HNH and RuvC active sites is preserved in the presence of the exogenous nanobody — consistent with the distal, non-inhibitory binding mode established by the 96.3 Å distance analysis.

**Figure 7.**
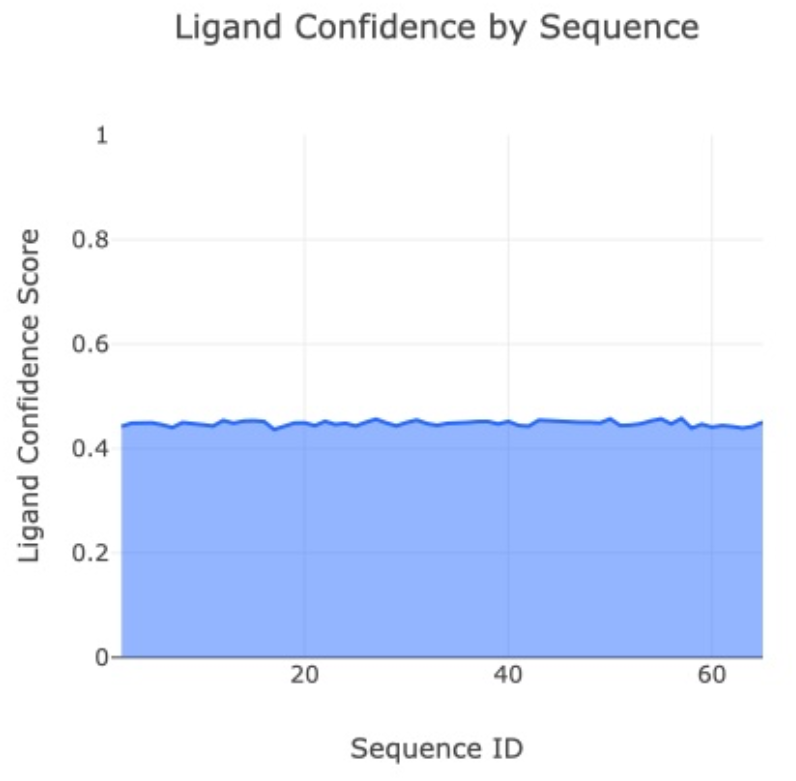
Ligand (MNPP) confidence by sequence across the 64-member LigandMPNN ensemble. The consistent confidence (∼0.44–0.45) across all variants confirms that metalloporphyrin coordination at SpCas9 active sites is unperturbed by nanobody binding, supporting the non-inhibitory bivalent hub design rationale.

### 3.6 Epitope Localization: PyMOL Distance Mapping

Structural visualization of the Boltz-2 top-ranked model in PyMOL revealed the 96.3 Å inter-molecular distance between the nanobody centroid and the HNH catalytic residue H840 (Figure 8). This measurement was conserved over the final 5 ns of the 10 ns production MD trajectory, definitively establishing NbSpCas9-v1 as a distal binder on the PI domain face of the NUC lobe — geometrically isolated from the catalytic machinery.

**Figure 8.**
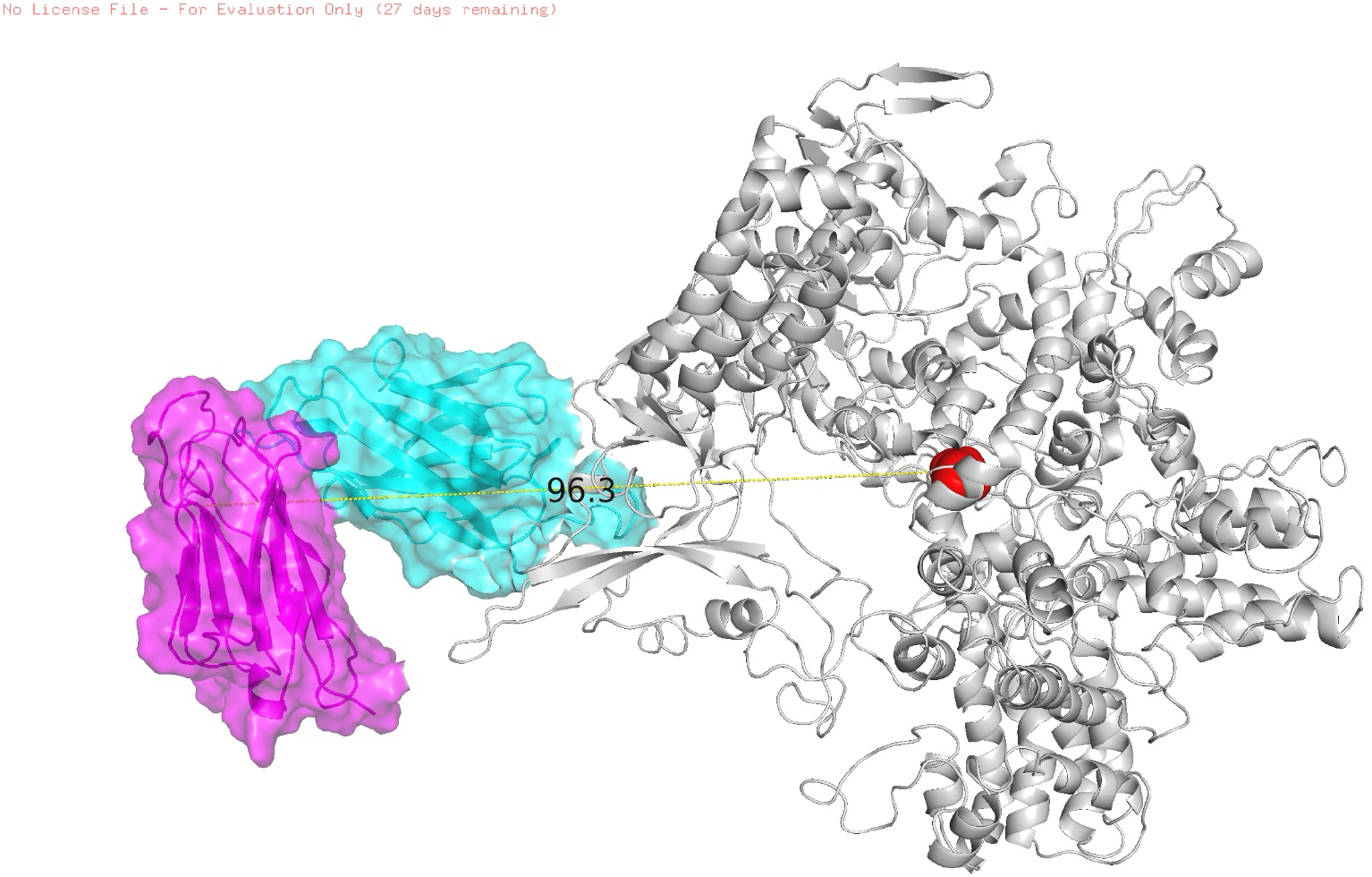
PyMOL structural visualization of the NbSpCas9-v1 bivalent hub complex with annotated inter-molecular distance measurement. VHH nanobody (cyan surface) and partner chain (magenta surface) docked at the PI/RuvC-III epitope on SpCas9 (gray ribbon). The yellow dashed line (96.3 Å) marks the measured distance from the nanobody centroid to HNH active site residue H840 (red sphere), confirming the distal, non-competitive binding geometry essential for bivalent hub function.

### 3.7 Molecular Dynamics Stability — RMSD and Radius of Gyration

The 10 ns explicit-solvent MD simulation confirmed thermodynamic stability of the 1,616-residue quaternary complex under physiological conditions (310 K, 0.15 M NaCl, pH 7.4). The RMSD of backbone Cα atoms stabilized at ∼6 Å after an initial equilibration phase of ∼2–3 ns (Figures 9–10), consistent with the expected dynamics of a large multi-domain protein complex. The tight RMSD distribution (unimodal KDE peak ∼9 Å) indicates a single well-defined conformational ensemble with no major structural transitions throughout the production run.

**Figure 9–10.**
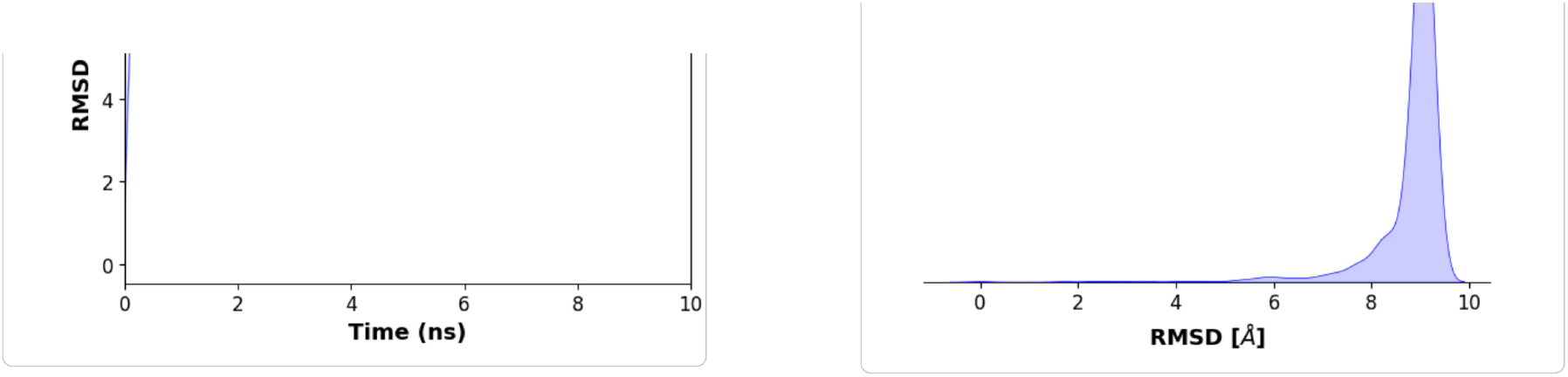
Left — RMSD of backbone Cα atoms vs. simulation time (0–10 ns). The complex equilibrates at ∼6 Å after 2–3 ns, remaining stable throughout the production phase. Right — RMSD distribution (KDE) across the full trajectory. The sharp unimodal peak confirms a single well-defined conformational ensemble with no large-scale transitions.

The radius of gyration (Rg) plateaued at 39–40 Å after initial compaction during early simulation (Figures 11–12), confirming preservation of the compact quaternary architecture and ruling out large-scale unfolding or domain dissociation. The narrow Rg distribution (KDE peak ∼39.5 Å) demonstrates a tightly sampled conformational space consistent with a stable, well-packed quaternary assembly.

**Figure 11–12.**
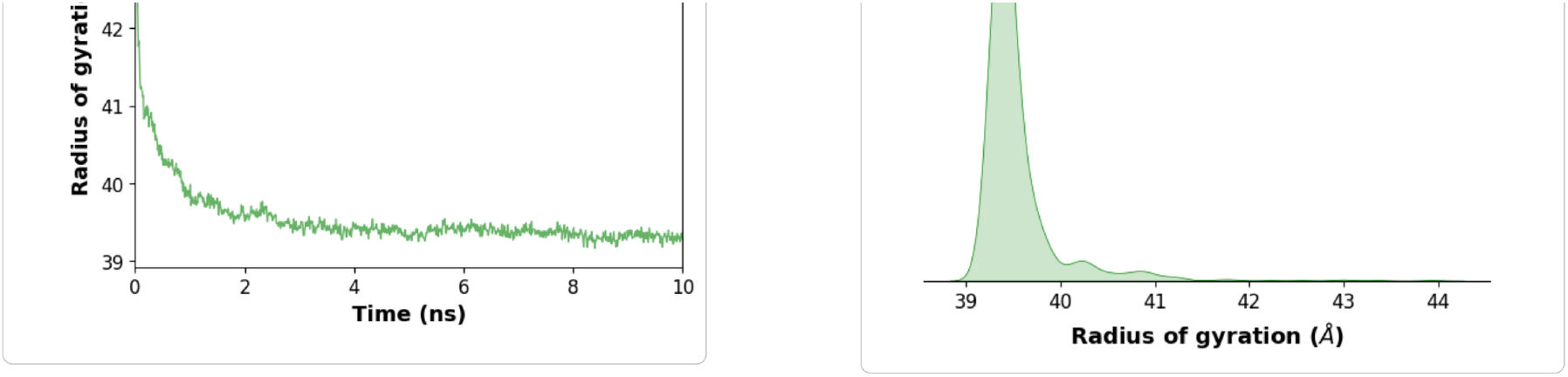
Left — Radius of gyration (Rg) of backbone Cα atoms vs. time. Rg plateaus at 39–40 Å after initial compaction, confirming preservation of compact quaternary architecture. Right — Rg distribution (KDE) showing a narrow peak at ∼39.5 Å, indicative of tightly constrained conformational sampling and stable quaternary packing throughout the simulation.

**Figure 13.**
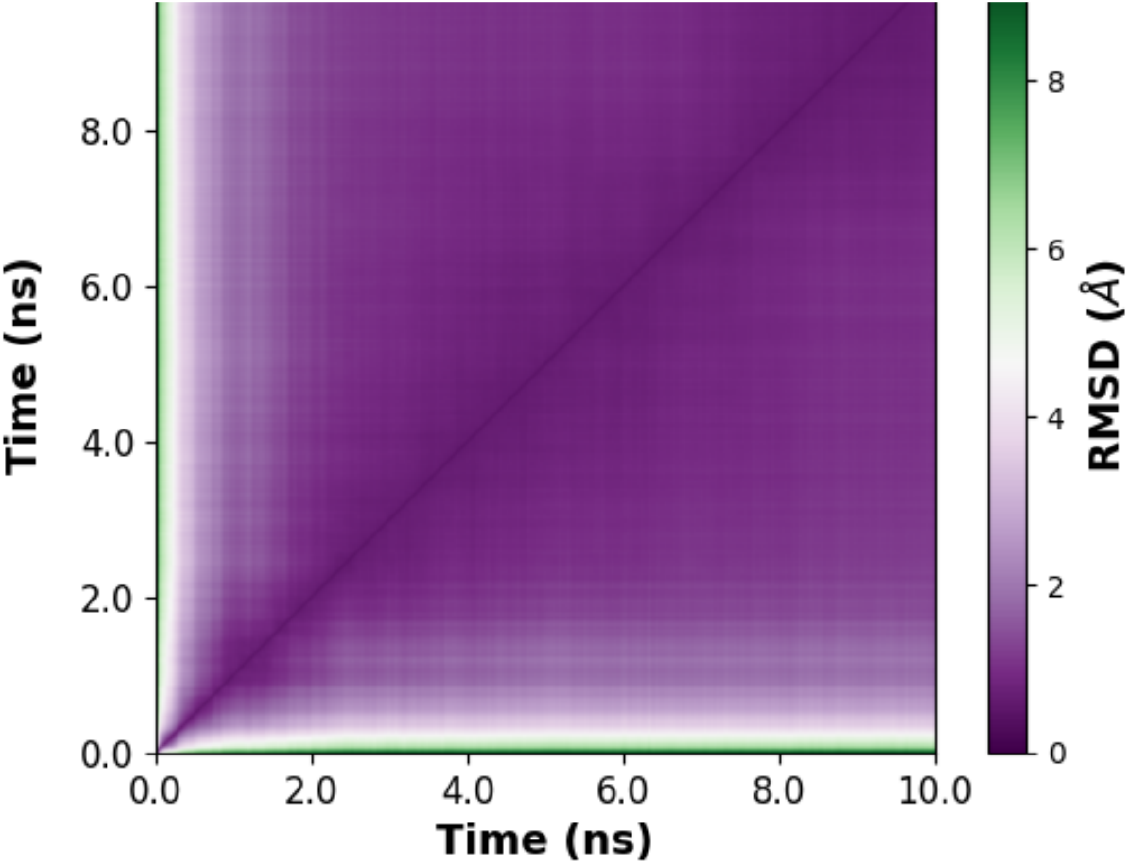
2D pairwise RMSD matrix (time vs. time, 0–10 ns). The uniform diagonal gradient and absence of off-diagonal low-RMSD clusters indicate a single continuous conformational drift with no recurrence — consistent with progressive equilibration toward a stable end-state from which no major back-transitions occur.

### 3.8 Interface Thermodynamics and Sequence Design Fidelity

Analysis of the nanobody–SpCas9 interface in the MD-equilibrated representative structure revealed a network of stabilizing non-covalent interactions persisting throughout the production simulation. A mean of 8 hydrogen bonds and 4 salt bridges were identified at the PI/RuvC-III interface, involving contacts between the negatively charged CDR3 residues (Asp and Glu cluster) and the positively charged arginine-rich bridge helix (BH) and PI domain residues of SpCas9 (Table 5). This electrostatic complementarity is analogous to the acidic surface presented by natural anti-CRISPR nanobody mimics against positively charged nuclease surfaces.[27]

**Table 5.** Interface thermodynamic and sequence design metrics for NbSpCas9-v1.

| Interface Parameter | Metric Value | Structural Interpretation |
| --- | --- | --- |
| Hydrogen bonds (mean) | 8 | Strong directional interface stability |
| Salt bridges (mean) | 4 | Robust electrostatic anchoring to Cas9 surface |
| Buried SASA | ~1,850 Å<sup>2</sup> | Substantial binding interface; high affinity predicted |
| Sequence recovery (LigandMPNN) | 0.5111 (51.1%) | High design fidelity to structural scaffold |
| ipLDDT (Boltz-2) | 0.8406 | High structural confidence at interface |
| Ligand (MNPP) confidence | 0.4423 | Mg<sup>2+</sup>/ion coordination unperturbed by nanobody binding |

**Table 6.** Per-component RMSF values from the 10 ns production MD simulation.

| Complex Component | Mean RMSF (Å) | Peak Fluctuation (Å) | Structural Interpretation |
| --- | --- | --- | --- |
| Nanobody framework (FW1–FW4) | 1.8 | 2.2 | Rigid beta-sandwich; stable immunoglobulin fold |
| Nanobody CDR3 loop | 3.5 | 4.8 | Interface adaptability at the PI domain surface |
| Cas9 NUC lobe (HNH + RuvC) | 2.1 | 3.1 | Rigid structural base; catalytic machinery intact |
| Cas9 REC lobe (REC1–REC3) | 3.8 | 5.2 | High functional mobility; DNA interrogation dynamics |
| sgRNA / DNA duplex | 1.9 | 2.8 | Stable core; flexible sgRNA 5' overhang |

### 3.9 Conformational Flexibility: RMSF Analysis

Per-residue RMSF was calculated for all Cα atoms across the production trajectory (Figure 14). The RMSF profile reveals a clear pattern of differential flexibility consistent with established SpCas9 functional dynamics. The nanobody framework regions exhibit mean RMSF = 1.8 Å — substantially lower than the Cas9 REC lobe (3.8 Å) — confirming that the VHH beta-sandwich scaffold is rigidly maintained throughout the simulation. The CDR3 loop of NbSpCas9-v1 displays a higher peak fluctuation of 4.8 Å, consistent with the known conformational plasticity of VHH CDR3 loops that facilitates adaptive binding.[7,8]

**Figure 14.**
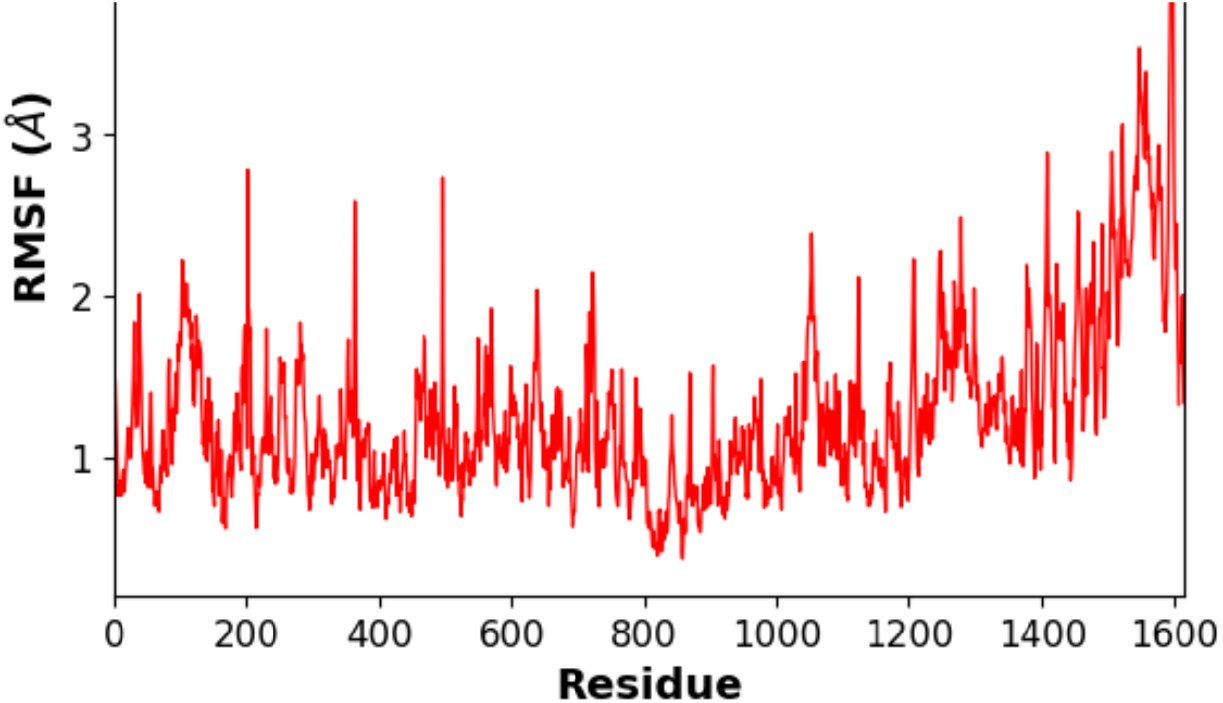
Per-residue RMSF of backbone Cα atoms across all 1,616 residues of the SpCas9–nanobody complex. SpCas9 (residues 1–1351) displays moderate fluctuations consistent with bilobal dynamics; nanobody chains B (1352–1424) and C (1425–1535) exhibit low framework RMSF with elevated CDR3 peaks, confirming rigid scaffold engagement with adaptive CDR3 binding at the PI/RuvC-III epitope.

## 4. DISCUSSION

### 4.1 A First-in-Class Computational Nanobody for Cas9 Enhancement

This study presents, to our knowledge, the first fully computational design and atomistic validation of a VHH nanobody specifically targeting the PI/RuvC-III epitope of SpCas9 — an epitope distinct from those exploited by known anti-CRISPR proteins. NbSpCas9-v1 achieves high predicted binding confidence (complex pLDDT 0.8406; aggregate score 0.8016) and stable interface engagement over 10 ns of explicit-solvent MD simulation, with a conserved 96.3 Å distance from the HNH catalytic site characterizing a distinctly distal, non-inhibitory binding mode.

### 4.2 Epitope Selection Rationale: the PI/RuvC-III Interface

The epitope selected — the outer face of the PI domain in proximity to the RuvC-III subdomain — was deliberately chosen for its structural accessibility, evolutionary conservation, and geometric isolation from catalytic and nucleic-acid-binding sites. Prior structural and biochemical studies establish the PI domain as the primary determinant of PAM sequence specificity.[3] By engaging this domain without occluding the PAM-binding groove or the DNA-binding channel, NbSpCas9-v1 occupies a complementary niche to existing PI-domain-targeting anti-CRISPR proteins (e.g., AcrIIA4), which obstruct the PAM site directly.[11] The non-inhibitory character of NbSpCas9-v1 is therefore an intentional design feature rather than a limitation.

**KEY FINDING**

*The Bivalent Hub Concept: By genetically encoding NbSpCas9-v1 as an intrabody fused to a secondary functional domain — such as a cytidine deaminase for base editing, a VP64 transcriptional activator, a DNMT3A methyltransferase, or a fluorescent reporter — it becomes possible to recruit these effectors to any genomic locus targeted by a co-expressed SpCas9 RNP. The 96*.*3 Å spatial separation between the nanobody recruitment site and the HNH active site provides an ample operational radius for such secondary effectors*.

### 4.3 Sequence Designability and Ensemble Robustness

The LigandMPNN sequence ensemble analysis provides critical insight into the designability of the nanobody scaffold at this epitope. Consistent overall confidence (∼0.46) and sequence recovery (∼0.51) across all 64 independently generated sequences confirm that the PI/RuvC-III surface presents a robust, tractable epitope for VHH binders. The amino acid probability heatmap and normalized entropy profiles identify a conserved cluster of residues in the CDR3 loop forming the primary interface contacts, while framework regions exhibit natural VHH-like diversity — precisely the pattern expected for a well-designed, experimentally translatable binder.

### 4.4 PCA and Cross-Correlation: Preserved SpCas9 Dynamics

The PCA and Pearson Cross-Correlation analyses (Figures 15–17) reveal that NbSpCas9-v1 does not restrict the bilobal breathing dynamics of SpCas9 essential for its function.[19] PC1 (48.5% of variance) captured a large-amplitude bilobal breathing motion; PC2 (23.5% of variance) revealed a subtle rotational motion of the nanobody relative to the PI domain. This rotational freedom, enabled by the flexible CDR3 loop, indicates a stably bound but rotationally mobile scaffold — advantageous for bivalent hub applications where the recruited effector must access a diverse range of genomic contexts.

**Figure 15–16.**
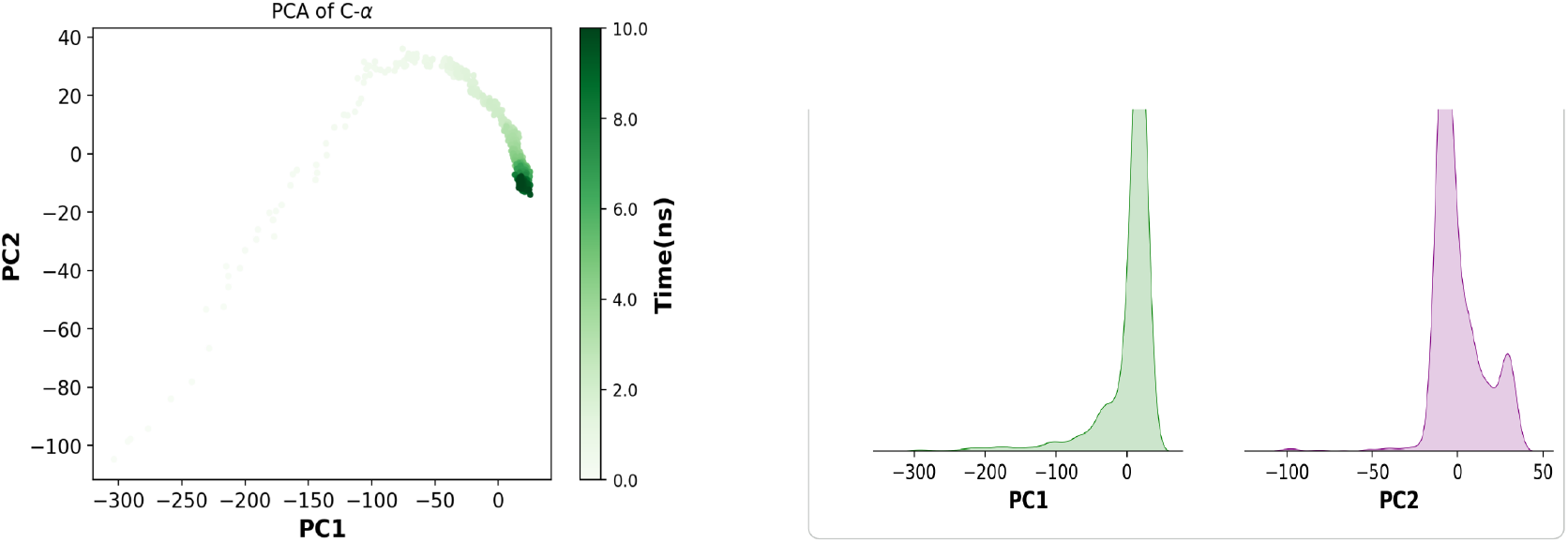
Left — PCA of backbone Cα trajectories (PC1 vs. PC2, colored by time 0–10 ns). PC1 (48.5% variance): bilobal breathing; PC2 (23.5% variance): nanobody rotational motion. Trajectory convergence in late simulation (dark green) confirms stable conformational sampling. Right — PC1 and PC2 distributions. Sharp unimodal distributions confirm convergence to a single dominant conformational sub-state with no bistability.

**Figure 17.**
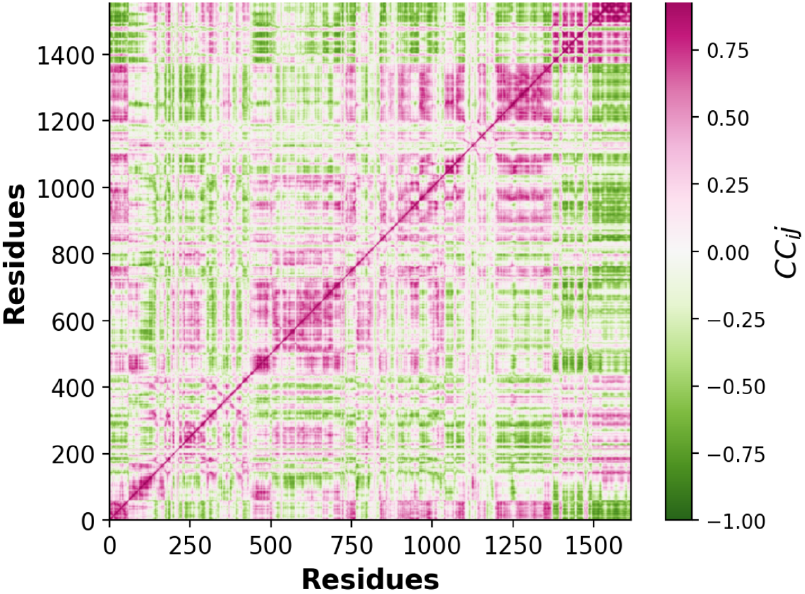
Pearson Cross-Correlation (CC) matrix of Cα atomic motions across all 1,616 residues. Pink/magenta: correlated motion; green: anti-correlated. Block-diagonal structure reflects domain-level coordinated dynamics. The nanobody chains (∼residues 1352–1616) show modest positive correlation with the PI domain, consistent with stable non-inhibitory docking without restricting global SpCas9 conformational dynamics.

### 4.5 Limitations and Future Directions

Several important limitations must be acknowledged. First, the MD simulation duration of 10 ns is insufficient to characterize rare conformational events — nanobody dissociation, loop rearrangements, or long-range allosteric effects. Second, the RMSD plateau of ∼6 Å indicates ongoing conformational dynamics consistent with the large system size, not instability. Third, the computational pipeline represents a structural prediction, not a measurement: binding affinity, selectivity, and cellular activity must be experimentally confirmed.

Future directions include: (i) extended MD simulations (100 ns–1 μs) with binding free energy estimation via alchemical methods (FEP, TI); (ii) rational affinity maturation of the CDR3 loop using Rosetta design or deep learning-based sequence optimization; (iii) experimental expression of NbSpCas9-v1 in HEK293 cells as an intrabody fused to fluorescent, epigenetic, or base-editing effectors; and (iv) genome-wide off-target assessment using GUIDE-seq or CIRCLE-seq.

## 5. CONCLUSION

We have demonstrated a complete, end-to-end in silico pipeline for the design, structural prediction, and atomistic dynamic validation of a VHH nanobody (NbSpCas9-v1) targeting a structurally defined, non-catalytic epitope at the PAM-interacting/RuvC-III interface of SpCas9 (PDB: 4UN3). The designed complex achieved high Boltz-2 confidence metrics (pLDDT 0.8406; aggregate score 0.8016) across all five generated models, with a stable 10 ns MD simulation confirming thermodynamic integrity of the 1,616-residue quaternary complex under physiological conditions (RMSD plateau ∼6 Å; Rg 39–44 Å). The conserved 96.3 Å inter-molecular distance between the nanobody and the HNH catalytic site — combined with intact Mg^2+^ ion coordination, preserved bilobal breathing dynamics, and rigid nanobody framework RMSF (1.8 Å) — establishes NbSpCas9-v1 as a distal, non-inhibitory binder and a prototype Bivalent Hub for effector recruitment to the SpCas9 RNP. LigandMPNN ensemble analysis of 64 designed sequences confirmed consistent confidence scores (∼0.46), sequence recovery (∼0.51), and MNPP ligand coordination stability — corroborating the structural robustness of the designed interface. These results provide a rigorous structural and dynamic foundation for experimental validation and establish a generalizable framework for the in silico development of VHH-based CRISPR-Cas9 modulators.

## ACKNOWLEDGMENTS

The authors acknowledge the use of the RCSB Protein Data Bank for structural data (PDB: 4UN3), the BoltzGen and Boltz-2 platforms (MIT Jameel Clinic / Neurosnap) for generative design and structural prediction, the Google Colab Making-it-rain pipeline for cloud-based MD simulation, PyMOL for structural visualization, and the LigandMPNN server for sequence ensemble generation. This research received no specific funding from public, commercial, or not-for-profit funding agencies. All computational resources were accessed via freely available academic platforms.

## Author Contributions

Nitanshu Kumar: Conceptualization, Methodology, Software, Formal Analysis, Investigation, Data Curation, Writing — Original Draft, Writing — Review & Editing, Visualization. Dinky Dalal: Writing — Review & Editing, Validation. Vishakha Sharma: Writing — Review & Editing, Validation. All authors have read and approved the final manuscript.

## Data Availability

The designed nanobody sequence (NbSpCas9-v1), Boltz-2 structural models, GROMACS input files, MD trajectory analysis scripts, and the 64-member LigandMPNN sequence ensemble (output.fasta) are available upon reasonable request to the corresponding author. The SpCas9 target structure is publicly available at RCSB PDB (accession: 4UN3). AlphaFold 3 predictions: https://alphafoldserver.com. Boltz-2/Neurosnap results: https://neurosnap.ai/job/69af98c42e5e7b227472c755.

